# From ΔΔ*G* Prediction to Mutation Prioritization with Conformal Prediction

**DOI:** 10.64898/2026.10.27.762132

**Authors:** Guido Barducci, Tiziana Sanavia, Piero Fariselli

## Abstract

Selecting mutations for protein engineering and studies of protein function requires deciding which predicted stability changes warrant experimental follow-up. A point estimate alone does not show whether uncertainty leaves both stabilizing and destabilizing effects plausible. To address this limitation, we introduce ProbJanusDDG, which combines a protein-stability predictor with standard and adaptive conformal prediction intervals. Calibration assigns equal total weight to each protein, accounting for uneven numbers of measured mutations.

For previously uncharacterized mutations, intervals lying entirely above or below zero provide a criterion for selecting candidate stabilizing or destabilizing substitutions, while intervals containing zero leave the direction unresolved. On the external low-homology S669L benchmark, adaptive intervals at the 50% nominal level assigned directional calls to 34.2% of mutations, with 94.8% observed sign accuracy. At the 70% level, 15.5% received calls, with 99.0% sign accuracy. At the 90% nominal level, standard and adaptive intervals achieved mutation-level coverage of 91.6% and 92.5%, whereas coverage averaged equally across proteins was approximately 87%.

Adaptive intervals redistributed width across mutations, with greater widening among predictions with larger observed errors and nearly unchanged mean width.

These results support uncertainty-aware screening of uncharacterized variants: prioritizing candidate stabilizing mutations for protein engineering and candidate destabilizing mutations for investigating stability-related effects on protein function. The method provides a selective criterion for experimental follow-up while distinguishing predictions with a resolved direction from those requiring further evidence.

**Significance statement:** Protein researchers often need to choose which mutations to test experimentally. We complement stability predictions with intervals that identify candidate stabilizing or destabilizing substitutions and leave uncertain directions unresolved. On external benchmarks, the selected subsets showed high observed sign accuracy. This provides an explicit criterion for prioritizing experiments in protein engineering and studies of stability-related functional effects.

## 1 Introduction

The change in folding free energy caused by a mutation, ΔΔ*G*, is a central quantity in protein engineering and in the interpretation of disease-associated variants[1][2][3].

An experimental study may seek substitutions that increase stability or variants that help test whether a functional change could be mediated by destabilization. In either setting, selecting mutations for characterization requires assessing the uncertainty of the predicted effect as well as its magnitude.

Numerous methods have been developed to predict ΔΔ*G*, which can be broadly divided into two main categories: structure-based and sequence-based approaches. Structure-based predictors exploit three-dimensional information about the protein, ranging from local atomic environments to residue- or atom-level interactions modeled through graph and geometric neural networks[4][5][6]. While these approaches can capture detailed structural context, they require an experimentally determined or predicted protein structure. Sequence-based methods, in contrast, operate directly on the amino-acid sequence and span a broad spectrum of complexity, from simple models based on handcrafted or one-hot encoded features to feed-forward neural networks and transformer-based protein language models, which provide contextual residue representations learned from large sequence collections[7][8]. Consequently, current protein-stability predictors differ substantially in their input information, representations, architectures, and overall model complexity.

Despite this diversity, most ΔΔ*G* predictors still provide only a point estimate, with no directly interpretable measure of the reliability of individual predictions[9][10]. Previous attempts to address this limitation include BayeStab, which uses a Bayesian neural network with concrete dropout to estimate predictive uncertainty and separate aleatoric and epistemic components [11]. However, its uncertainty estimates are primarily used to characterize sources of prediction error and are not calibrated against prespecified coverage levels. Consequently, they do not directly provide a prediction interval with an interpretable reliability level for an individual mutation.

A complementary approach was proposed by Sapozhnikov et al. [12], who modeled the absolute error of FoldX predictions using linear regression and constructed mutation-specific prediction intervals. Although these intervals support uncertainty-aware interpretation of stability effects, their validity relies on the assumptions underlying the regression model, rather than distribution-free conformal calibration. Moreover, folding-stability validation involved only ten proteins, and aggregate coverage was calculated by pooling mutations, giving greater influence to proteins with more measurements. Despite approximately 95% aggregate coverage, coverage for individual proteins fell as low as 79%. These results highlight the importance of distinguishing mutation-level coverage from coverage averaged equally across proteins when assessing generalization to previously unseen proteins.

Moreover, the required level of information depends on the application: in some settings an accurate estimate of the magnitude of ΔΔ*G* is important, whereas in others it may be sufficient to determine, with quantified uncertainty, whether a mutation is expected to stabilize or destabilize the protein [13]. Conformal prediction intervals provide a natural way to address both settings, by quantifying uncertainty around the predicted magnitude and by distinguishing predictions whose plausible values lie entirely on one side of zero from those for which the direction of the effect remains uncertain. These considerations motivate uncertainty-quantification strategies that are both reliable and largely independent of the underlying predictive architecture[14][15][16].

In this paper, we introduce a protein-aware conformal approach for uncertainty-aware ΔΔ*G* prediction, instantiated in ProbJanusDDG, a predictor that provides both point estimates and mutation-specific uncertainty estimates. The model is designed to produce an exactly antisymmetric mean prediction together with an exactly symmetric mutation-specific uncertainty estimate. This scale is used within a protein-weighted CV+ procedure to construct both standard and adaptive conformal prediction intervals, with adaptive widths modulated according to the predicted relative error scale. Calibration is based on held-out residuals and predictions from the corresponding fold model, while equal total weight is assigned to each WT protein to limit the influence of proteins represented by many mutations. Point predictions are obtained as the median across five fold models. The overall prediction and calibration procedure, including ensemble aggregation across fold models, is illustrated in Figure 4b.

Although developed here in conjunction with ProbJanusDDG, the conformal calibration strategy is model-agnostic and can, in principle, be applied to predictors with different architectures and input representations. To illustrate this property in practice, we additionally apply the same calibration procedure to a deliberately simple model based on one-hot encoded sequence features. This provides a contrasting test case with substantially different representation, architecture, and predictive performance, while leaving the calibration strategy unchanged.

## 2 Results

### 2.1 Evaluation strategy

We evaluated the proposed approach in three complementary steps. First, we assessed whether the probabilistic formulation and ensemble procedure (see Section 4.3) preserved competitive ΔΔ*G* point-prediction performance on the external low-homology benchmarks. We then examined whether the learned mutation-specific scale carried information about prediction error and whether it could be used to construct well-calibrated standard and adaptive conformal prediction intervals. Finally, we investigated the practical behavior of these intervals, including their width allocation, their ability to provide stabilizing or destabilizing directional calls, and the transfer of the calibration procedure to a deliberately simple one-hot baseline model.

This evaluation separates the quality of the underlying point predictor from the quality of the uncertainty quantification: accurate predictions are desirable, but the central objective is to determine whether individual predictions can be accompanied by informative and empirically calibrated uncertainty estimates.

#### 2.2 Nested epoch selection and point-prediction performance

Before evaluating uncertainty quantification through conformal prediction, we first verified that the probabilistic training and ensemble procedure preserved the point-prediction accuracy of the original JanusDDG model[7] and remained competitive with existing state-of-the-art methods.

The internal validation procedure selected different training durations across the five outer folds, with optimal epochs of 57, 232, 126, 283, and 105 for folds 0–4, respectively (Supplementary Figure 7). This variability is expected because each outer-fold model is associated with a different three-fold training subset and a different set of proteins used for internal validation. Importantly, the corresponding outer fold was never used for epoch selection, ensuring that its residuals remained fully out-of-sample with respect to both parameter fitting and model selection.

After the optimal epoch had been identified for each split, the corresponding model was reinitialized and refitted on all four non-outer folds for the selected number of epochs. All five four-fold refits were completed before any conformal prediction analysis was performed. Point predictions were obtained as the median of the predictions produced by these five refitted models.

We then evaluated the resulting median ensemble on the external S669L and S461L benchmarks using Pearson and Spearman correlation, mean absolute error (MAE), and mean squared error (MSE). On S669L, ProbJanusDDG achieved a Pearson correlation of *r* = 0.53 and an MAE of 0.99 kcal mol^−1^, closely matching the original JanusDDG results (*r* = 0.55, MAE = 0.97 kcal mol^−1^). On S461L, it achieved *r* = 0.70 and an MAE of 0.74 kcal mol^−1^, again comparable to JanusDDG (*r* = 0.69, MAE = 0.71 kcal mol^−1^). The complete performance metrics and comparison with previously reported methods are provided in Supplementary Tables 3 and 4.

Thus, introducing the training and ensemble framework required for the subsequent uncertainty analysis did not result in a meaningful loss of predictive accuracy. Having established that the underlying point predictor retained performance comparable to JanusDDG and other competitive protein-stability predictors, we next assessed whether its predictions could be complemented by statistically calibrated uncertainty estimates using conformal prediction.

#### 2.2.1 Predicted uncertainty tracks prediction error

We first investigated whether the model-derived uncertainty scale was associated with the magnitude of the prediction error. Specifically, we computed the Spearman correlation between the mean fold-specific scale percentile and the absolute error of the median prediction. The two quantities were positively associated, with Spearman *ρ* = 0.22 on S669L and *ρ* = 0.23 on S461L. These modest correlations indicate that the learned scale contains information about the relative difficulty of individual predictions.

Although the scale should not be interpreted as a precise estimate of the error for an individual mutation, this association is consistent with its intended role as an indicator of prediction difficulty: mutations with larger observed errors tended to show larger increases in interval width relative to standard CV+ intervals (Fig. 1C). Thus, the predicted scale provides a relative uncertainty signal that can be used to adapt interval widths across mutations.

**Figure 1:**
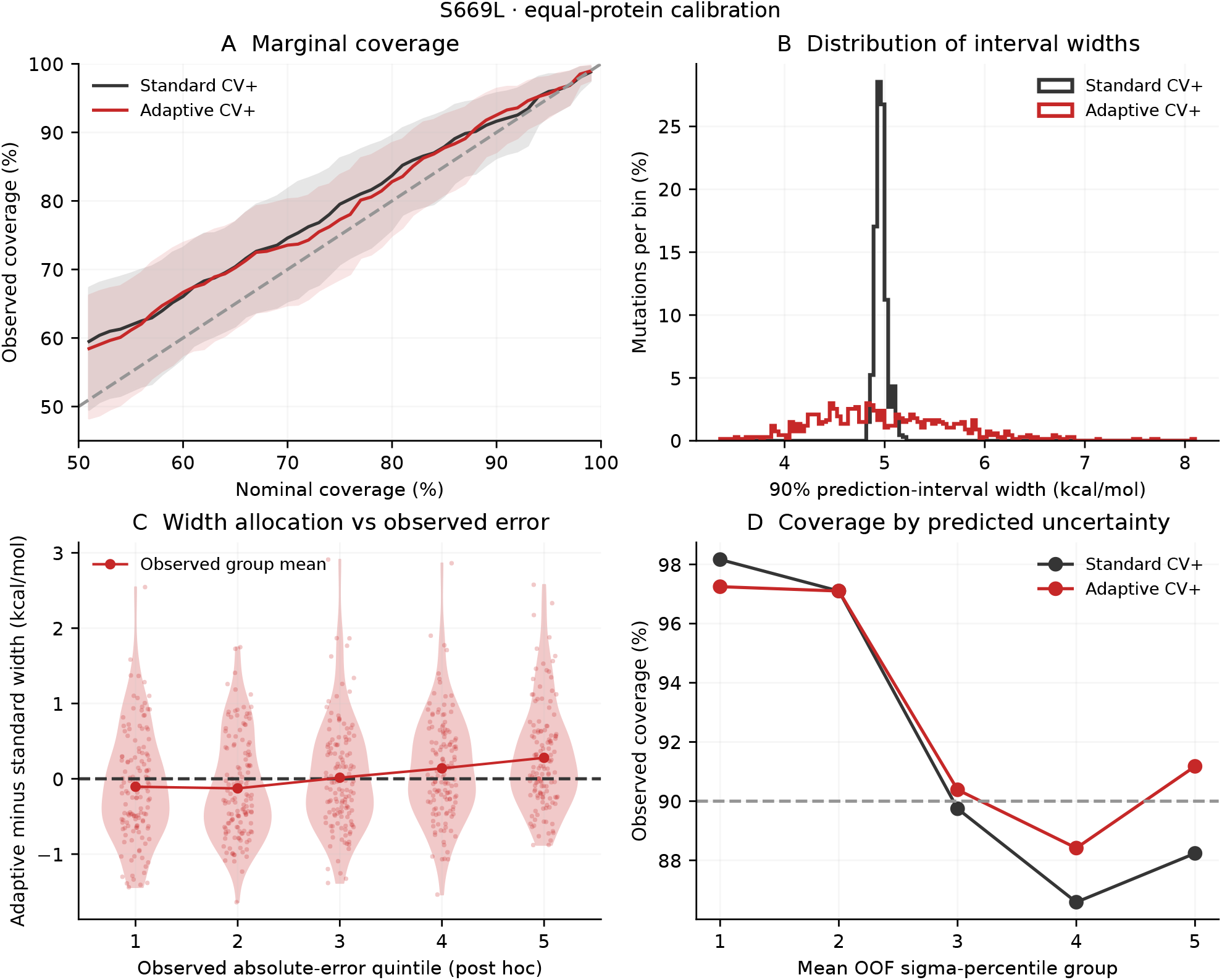
What does predicted scale add to standard CV+ intervals? S669L results use the five nested-MSE refits, median point prediction and empirical protein-weighted calibration from S2450. Black denotes standard and red adaptive CV+. (A) Mutation-weighted observed versus nominal coverage at 49 levels (51–99%), with pointwise 95% protein-cluster bootstrap bands (4,000 resamples). (B) Distributions of both interval widths at nominal 90%; standard CV+ is not exactly constant-width. (C) Adaptive-minus-standard width in post-hoc quintiles of observed absolute median-prediction error: violins and small points show mutations, connected large points show group means. (D) Mutation-weighted coverage at 90% within fixed uncertainty-percentile groups. Groups in C are not used for calibration. The weighting of plotted external observations is distinct from the equal-protein weighting used to construct intervals.

#### 2.2.2 Marginal and protein-averaged coverage

Empirical mutation-level coverage was defined as the percentage of test mutations whose observed ΔΔ*G* fell within the prediction interval, with each mutation contributing equally. To summarize agreement with nominal coverage, we calculated the mean absolute difference between empirical and nominal coverage across the 49 integer nominal levels from 51% to 99%. Smaller values indicate closer agreement; the absolute difference does not distinguish undercoverage from overcoverage.

On S669L, this mean absolute difference was 4.06 percentage points for standard CV+ and 3.63 for adaptive CV+ (Fig. 1A). On S461L, the corresponding values were 11.90 and 11.63 percentage points (Fig. 5), mainly reflecting coverage above the nominal levels.

The greater overcoverage on S461L is consistent with its lower prediction errors. Both benchmarks used the same calibration residuals from S2450, and mean interval widths at the 90% nominal level were approximately 5 kcal mol^−1^ on both datasets. Smaller errors with similarly wide intervals can produce higher coverage, explaining why better point-prediction accuracy can coexist with larger deviations from nominal coverage. Since S461L is a curated subset of S669L rather than a random subsample, these results also suggest that empirical coverage depends on benchmark composition.

At 90% nominal coverage on S669L, pooling mutations gave approximately 92% coverage, whereas averaging equally across proteins gave approximately 87% (Table 1). This distinction matters when assessing predictions for a new protein, rather than predominantly for proteins with many measured mutations.

**Table 1.** Empirical performance of protein-weighted CV+ intervals at nominal 90%. Coverage is reported both per mutation and averaged equally over WT proteins. Width and interval score (IS) are mutation-weighted and in kcal mol^−1^; dispersion is the population SD of mutation-weighted coverage across five predicted-uncertainty groups, in percentage points.

| Dataset | Interval | Mutation coverage (%) | Protein coverage (%) | Mean width | IS | Dispersion |
| --- | --- | --- | --- | --- | --- | --- |
| S669L | Standard | 91.63 | 86.76 | 4.964 | 6.716 | 4.75 |
|  | Adaptive | 92.53 | 86.87 | 5.004 | 6.551 | 3.63 |
| S461L | Standard | 97.61 | 96.56 | 4.965 | 5.188 | 2.71 |
|  | Adaptive | 98.05 | 93.85 | 4.967 | 5.109 | 1.47 |

#### 2.2.3 Behavior of adaptive prediction-interval widths

Analysis of prediction-interval widths showed that adaptive CV+ produced mutation-specific intervals with substantially greater width variability than standard CV+.

Standard CV+ produced intervals with only limited variation in width across S669L, ranging from 4.82 to 5.19 kcal mol^−1^. In contrast, adaptive CV+ generated substantially more heterogeneous intervals, with widths ranging from 3.36 to 8.10 kcal mol^−1^ (Fig. 1B). Relative to the standard intervals, 352 of the 669 adaptive intervals became narrower, whereas 317 became wider. Despite this marked redistribution at the level of individual mutations, the average interval width remained essentially unchanged, increasing only from 4.96 to 5.00 kcal mol^−1^. These results indicate that the adaptive procedure primarily reallocates interval width across mutations rather than systematically increasing or decreasing the overall width.

In the error-stratified analysis, the mean adaptive-minus-standard interval width was −0.11 kcal in the lowest absolute-error quintile and +0.28 kcal in the highest (Fig. 1C), corresponding to a high-minus-low contrast of 0.39 kcal. On S461L, the corresponding contrast was 0.427 kcal. These results indicate an association between larger observed prediction errors and greater adaptive widening relative to standard CV+.

To illustrate the practical difference between standard and adaptive intervals at the 90% nominal level, we examined the mutations whose observed ΔΔ*G* values changed coverage status between the two interval definitions. Adaptive intervals covered seven S669L mutations missed by standard CV+, while one previously covered mutation became uncovered. On S461L the counts were four and two, respectively. Figure 2 shows all S669L switches, including the loss, with their actual interval endpoints. Width-stratified examples selected without reference to observed error are shown in Fig. 6.

**Figure 2:**
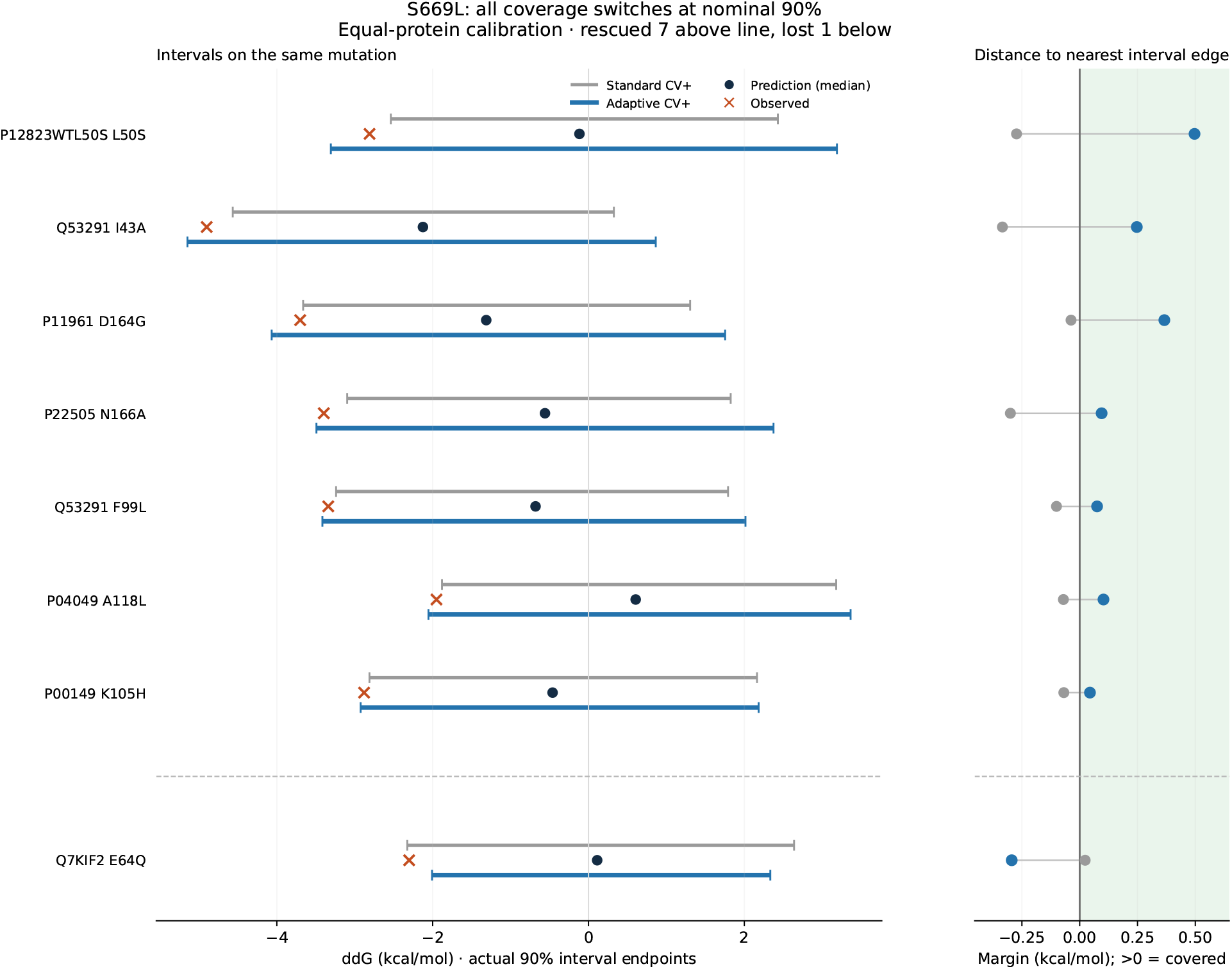
All S669L coverage switches between the standard and adaptive protein-weighted 90% intervals, using median point predictions. Seven rescued mutations are above the dashed line and the single lost mutation is below. Left: true interval endpoints (gray standard, blue adaptive), shared median prediction and measured target. Right: the signed margin min(*y* − *L, U* − *y*), positive when covered and negative when missed. The cases are selected post hoc by their coverage outcome and are illustrative rather than a performance estimate.

### 2.2.4 Interval-based stabilizing and destabilizing calls

To quantify how often the predictive intervals identify the direction of a mutation-induced stability change, we evaluated each integer nominal level *c* from 50% to 95%, inclusive. For each level, we recomputed the adaptive empirical CV+ endpoints using the existing OOF residuals and equal-protein calibration weights, with *α* = 1 −*c/*100. The model checkpoints and median point predictor were fixed. Evaluation used the original benchmark mutations without adding generated reverses, so each experimentally characterized mutation contributed once.

Under our sign convention, a mutation is called stabilizing when the entire interval lies strictly above zero (*L*_*i*_(*c*) *>* 0), and destabilizing when it lies strictly below zero (*U*_*i*_(*c*) *<* 0). If the interval contains or touches zero, its direction is considered indeterminate at that nominal level. Thus the number of direction calls is

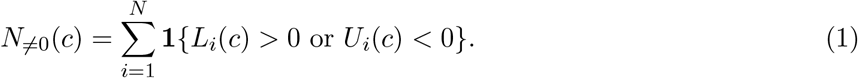

The mutation-level call rate is 100*N*_≠0_(*c*)*/N*, using every mutation in the test set as the denominator, including indeterminate cases.

Because several mutations belong to the same protein, we also count the distinct full-WT protein sequences with at least one direction call:

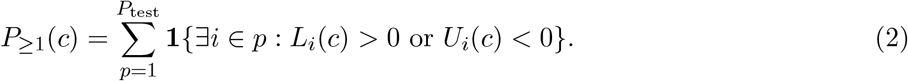

Their percentage is 100*P*≥_1_(*c*)*/P*_test_, with *P*_test_ = 87 for S669L and 46 for S461L. Each protein is counted once even if it contains multiple classified mutations or both stabilizing and destabilizing calls. This statistic does not assert that all mutations of that protein have a determined direction. Protein identifiers that map to the same full WT sequence are grouped together, consistently with calibration.

Figure 3 summarizes selected nominal levels and Supplementary Table 5 provides a more complete summary of the results across nominal coverage levels ranging from 50% to 90%. We highlight the 50%, 70%, and 90% nominal levels to illustrate the relationship between call rate and observed sign accuracy. At 50%, 229 of 669 S669L mutations (34.23%) and 190 of 461 S461L mutations (41.21%) exclude zero, spanning 46 of 87 proteins (52.87%) and 26 of 46 proteins (56.52%), respectively. Observed sign accuracy among these calls was 94.76% (217/229) on S669L and 98.42% (187/190) on S461L. At 70%, the call rates decreased to 15.55% (104/669) and 20.82% (96/461), while sign accuracy increased to 99.04% (103/104) and 100% (96/96), respectively. At 90%, these counts fall to four mutations (0.60%) from two proteins (2.30%) on S669L, and five mutations (1.08%) from two proteins (4.35%) on S461L; all of these 90% calls are destabilizing in the original benchmark orientation and have correct signs. As the nominal level increases, the intervals expand and the set of direction calls can only shrink, a nesting property also checked numerically.

**Figure 3:**
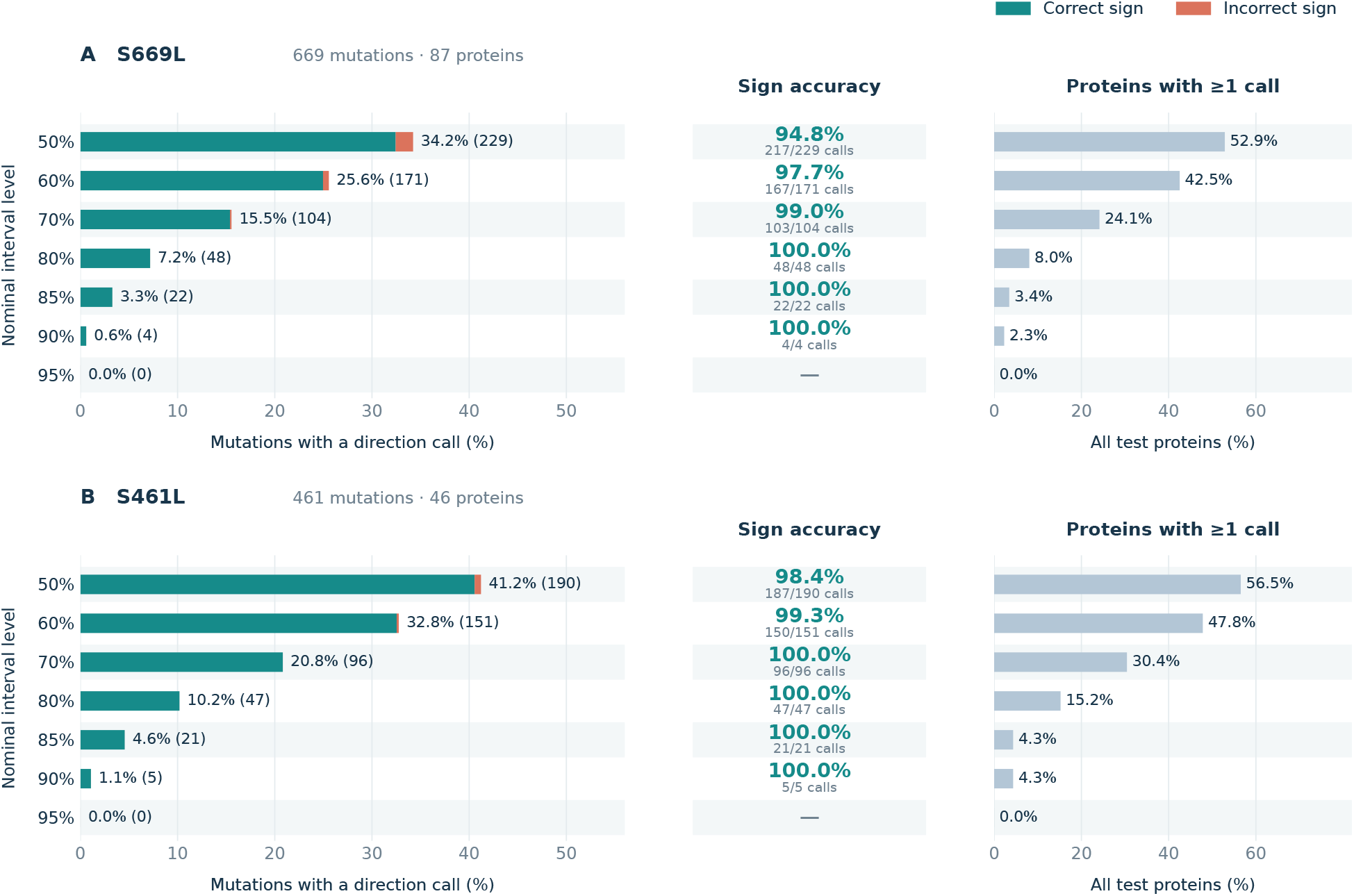
Direction calls and observed sign accuracy for S669L (A) and S461L (B) at selected nominal interval levels. Results use the original benchmark mutation directions, without generated reverses. Calls are assigned when adaptive protein-weighted CV+ intervals lie strictly above or below zero. Left: percentage of all test mutations receiving a call, partitioned into correct and incorrect signs; labels give percentages and counts in parentheses. Center: observed sign accuracy among called mutations, with correct/total counts. Right: percentage of distinct full-WT proteins with at least one call. Calibration weights proteins equally, whereas sign accuracy weights called mutations equally. Observed zero targets count as incorrect; a dash indicates no calls. Empirical sign accuracy is not a nominal coverage guarantee. S461L overlaps S669L.

Most calls were destabilizing in the original benchmark orientation. Reversing a mutation changes the sign of the model’s predicted ΔΔ*G* while preserving its uncertainty scale. For a reflected interval [−*U*_*i*_(*c*), −*L*_*i*_(*c*)], the directional call is likewise reversed, with unchanged correctness against the sign-inverted target.

For many practical applications, distinguishing whether a mutation is stabilizing or destabilizing can be more relevant than precisely estimating the magnitude of its ΔΔ*G*. For this reason, we additionally report observed sign accuracy among the zero-excluding calls:

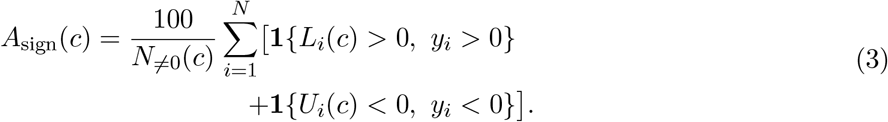

Each called mutation contributes equally to this accuracy, independently of the protein weights used for calibration. An observed target equal to zero does not count as a correct directional call. Accuracy is undefined (shown as –) when no interval excludes zero. For an uncharacterized mutation, the same rule assigns a call using only the interval endpoints; the observed target is used only when evaluating sign accuracy. Call rates and empirical sign accuracy are distinct from observed interval coverage and from probabilities that individual assigned signs are correct. In particular, a 90% nominal interval does not establish 90% accuracy among the selected mutations; such a selected-subset guarantee is not supplied by this empirical protein-weighted construction.

These results support zero-exclusion as a practical criterion for selecting a subset of mutations for experimental stability measurements. The call rate indicates the fraction of candidates retained, while observed sign accuracy describes the correctness of their predicted directions within that subset.

#### 2.2.5 Calibration with the baseline model

To assess whether the calibration procedure transfers to a substantially different predictor, we evaluated the simple sequence-based baseline described in Materials and methods, which uses full-length one-hot encoded WT and mutant sequences without pretrained embeddings. In this comparison, both the input representation and model architecture are changed, while the training, validation, aggregation, and conformal calibration procedures are kept unchanged.

Point-prediction performance was lower than that of the main model. On S669L, the median baseline prediction achieved Pearson *r* = 0.33, Spearman *ρ* = 0.31, MAE 1.23 kcal mol^−1^, and MSE 2.76 (kcal mol^−1^)^2^. On S461L, the corresponding values were *r* = 0.43, *ρ* = 0.40, MAE 1.00 kcal mol^−1^, and MSE 1.74 (kcal mol^−1^)^2^. Nevertheless, at the nominal 90% level, standard and adaptive intervals covered 90.88% and 90.28% of S669L mutations, respectively (Table 2). Their mean widths were 5.64 and 5.84 kcal mol^−1^, compared with 4.96 and 5.00 kcal mol^−1^ for ProbJanusDDG. Coverage averaged equally across S669L proteins was lower, at 86.31% and 86.41%. On S461L, mutation-level coverage was conservative for both interval rules, while protein-averaged coverage was 91.59% and 92.00%.

**Table 2.** One-hot baseline intervals at nominal 90% coverage, using equal total calibration weight per WT protein. Mutation coverage weights test mutations equally; protein coverage averages within each WT protein and then equally across proteins. Mean width weights test mutations equally and is expressed in kcal mol^−1^. S461L overlaps S669L.

| Dataset | Interval | Mutation | Protein | Width |
| --- | --- | --- | --- | --- |
| S669L | Standard | 90.88% | 86.31% | 5.64 |
|  | Adaptive | 90.28% | 86.41% | 5.84 |
| S461L | Standard | 95.88% | 91.59% | 5.64 |
|  | Adaptive | 96.31% | 92.00% | 5.93 |

**Table 3.** Point-prediction performance of the median ensemble on the external test sets.

| Dataset | Pearson $r$ | Spearman $\rho$ | MAE | MSE |
| --- | --- | --- | --- | --- |
| S669L | 0.53 | 0.55 | 0.99 | 2.03 |
| S461L | 0.70 | 0.68 | 0.74 | 0.97 |

**Table 4.**
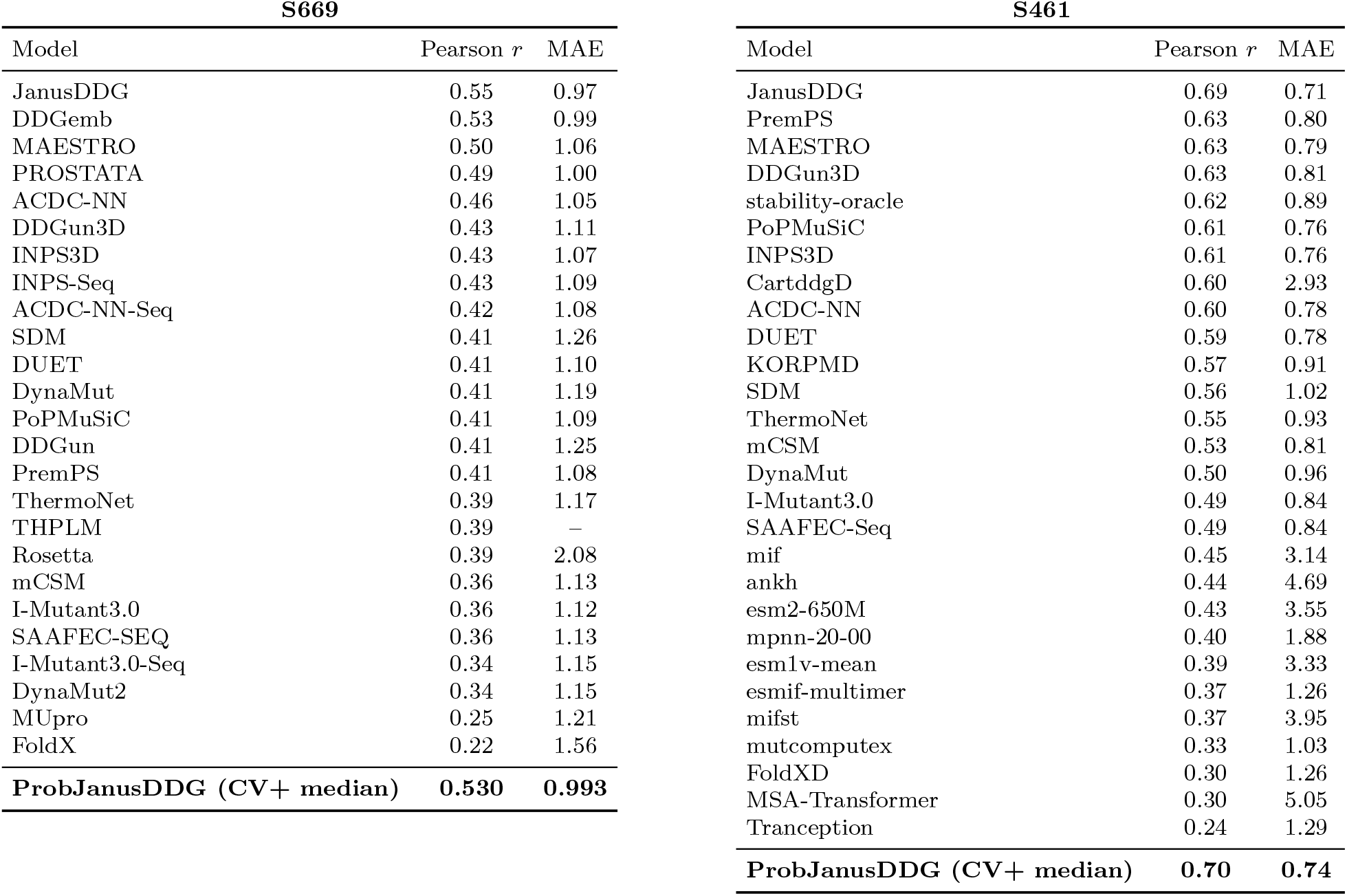
Point-prediction performance on the external S669 and S461 benchmarks. Comparative values and the original JanusDDG results are taken from the source data of Fig. 3 of the JanusDDG paper. ProbJanusDDG denotes the median of the five models selected by protein-weighted MSE and refitted on four folds. Its rows use the local S669L and S461L datasets. MAE is reported in kcal mol^−1^. The S461 values refer to the direct-mutation evaluation.

**Table 5.** Direction calls from adaptive protein-weighted CV+ intervals at each nominal level. *N*_+_ and *N*_−_ count intervals strictly above and strictly below zero. *N*_=0_ is their sum; its percentage uses all test mutations (669 or 461). *P*≥_1_ counts distinct WT proteins with at least one such mutation; its percentage uses all test proteins (87 or 46). Proteins can have both types of mutation and are counted once in *P*≥_1_. Sign acc. is the percentage of zero-excluding calls whose assigned sign matches the observed target, with equal weight per called mutation. Zero targets count as incorrect; – denotes no calls. This empirical accuracy is distinct from nominal coverage.

| S669L: 669 mutations, 87 proteins |  |  |  |  |  | S461L: 461 mutations, 46 proteins |  |  |  |  |  |
| --- | --- | --- | --- | --- | --- | --- | --- | --- | --- | --- | --- |
| Nominal (%) | $N_+$ | $N_-$ | $N_{\neq 0}$ (%) | $P_{\geq 1}$ (%) | Sign acc. (%) | $N_+$ | $N_-$ | $N_{\neq 0}$ (%) | $P_{\geq 1}$ (%) | Sign acc. (%) | |
| 50 | 10 | 219 | 229 (34.23) | 46 (52.87) | 94.76 | 7 | 183 | 190 (41.21) | 26 (56.52) | 98.42 |  |
| 55 | 6 | 195 | 201 (30.04) | 41 (47.13) | 96.52 | 4 | 167 | 171 (37.09) | 23 (50.00) | 98.83 |  |
| 60 | 3 | 168 | 171 (25.56) | 37 (42.53) | 97.66 | 3 | 148 | 151 (32.75) | 22 (47.83) | 99.34 |  |
| 65 | 2 | 135 | 137 (20.48) | 29 (33.33) | 98.54 | 2 | 120 | 122 (26.46) | 18 (39.13) | 100.00 |  |
| 70 | 1 | 103 | 104 (15.55) | 21 (24.14) | 99.04 | 1 | 95 | 96 (20.82) | 14 (30.43) | 100.00 |  |
| 75 | 1 | 73 | 74 (11.06) | 13 (14.94) | 100.00 | 1 | 70 | 71 (15.40) | 11 (23.91) | 100.00 |  |
| 80 | 0 | 48 | 48 (7.17) | 7 (8.05) | 100.00 | 1 | 46 | 47 (10.20) | 7 (15.22) | 100.00 |  |
| 85 | 0 | 22 | 22 (3.29) | 3 (3.45) | 100.00 | 0 | 21 | 21 (4.56) | 2 (4.35) | 100.00 |  |
| 90 | 0 | 4 | 4 (0.60) | 2 (2.30) | 100.00 | 0 | 5 | 5 (1.08) | 2 (4.35) | 100.00 |  |

These results support the practical applicability of the calibration procedure across contrasting predictors: lower point accuracy can coexist with similar mutation-level coverage, at the cost of wider intervals. The comparison therefore separates the empirical calibration of intervals from the predictive information that determines their usefulness and width.

## 3 Discussion and Conclusions

The practical contribution of this study is to complement ΔΔ*G* estimates with an explicit criterion for choosing mutations for experimental characterization. Prediction intervals allow researchers to identify candidate substitutions whose predicted effects remain stabilizing or destabilizing across the interval, and to leave the direction unresolved for others. The protein-weighted CV+ construction accounts for uneven numbers of measured mutations across proteins, while ProbJanusDDG supplies competitive point predictions and mutation-specific relative uncertainty estimates. Together, these components connect uncertainty quantification to decisions about which variants to investigate.

Previous work on uncertainty quantification in protein-stability prediction includes BayeStab [11]. BayeStab incorporated Bayesian uncertainty estimation into protein-stability prediction and examined contributions from model and data uncertainty. However, that study did not report empirical interval coverage at prespecified nominal levels. The present work provides a practical basis for experimental mutation selection by calibrating mutation-specific prediction intervals using held-out errors and assessing their empirical coverage, while accounting for uneven numbers of measured mutations across proteins. For mutations without experimental measurements, intervals lying entirely above or below zero provide an explicit criterion for prioritizing candidate stabilizing or destabilizing substitutions. On the external benchmarks, this criterion identified subsets with high observed sign accuracy (Fig. 3). Increasing the nominal interval level makes selection more stringent, allowing researchers to balance the strength of the selection criterion against the number of candidates retained for experiments. The same calibration strategy could, in principle, also be applied to all stability predictors, including BayeStab.

Within ProbJanusDDG, the learned scale provides information about relative prediction reliability. Its association with absolute prediction error is modest, indicating that *σ* should not be interpreted as an accurate estimate of the error of an individual mutation. Rather, it acts as a relative uncertainty signal that can be used to adapt interval widths across predictions. In the conformal construction, the predicted scale modulates residual normalization and therefore the relative allocation of interval width, whereas the final interval endpoints are determined empirically from held-out residuals.

The probabilistic formulation also preserves the thermodynamic symmetry expected under mutation reversal. While the predicted ΔΔ*G* changes sign when the mutation direction is reversed, the associated uncertainty remains unchanged. Enforcing these properties algebraically guarantees consistency between forward and reverse predictions and prevents contradictory uncertainty estimates for the same mutation pair.

A second important result concerns the evaluation of predictive uncertainty. Similar marginal coverage can mask substantial differences in how uncertainty is distributed across mutations and proteins. In our experiments, standard and adaptive CV+ intervals achieved comparable overall coverage, but adaptive intervals redistributed width more strongly across individual predictions. This redistribution was associated with observed prediction error, with greater widening among mutations with larger errors, while the average interval width remained nearly unchanged. Adaptive intervals also reduced coverage dispersion across predicted-uncertainty groups. At the same time, improved mutation-level coverage did not necessarily translate into improved protein-averaged coverage. These observations support evaluating conformal intervals using complementary criteria, including mutation-level coverage, protein-averaged coverage, interval width, and coverage heterogeneity.

The directional-call analysis provides a direct practical interpretation of the resulting intervals. When an interval lies entirely above or below zero, the predicted effect can be assigned a stabilizing or destabilizing direction; when the interval contains zero, the method abstains from assigning a direction at that nominal level. On S669L, this criterion yielded 94.8% observed sign accuracy for 34.2% of mutations at the 50% nominal interval level, and 99.0% accuracy for 15.5% of mutations at the 70% level (Fig. 3). Thus, increasing the nominal level produces a more selective set of directional calls, trading the number of classified mutations for higher observed sign accuracy. These values are empirical accuracies within the selected subsets and should not be interpreted as guarantees implied by the nominal interval level.

In a prospective protein-engineering screen, intervals entirely above zero could be used to select candidate stabilizing substitutions for experimental characterization. Intervals entirely below zero could similarly identify candidate destabilizing variants for testing whether changes in stability contribute to altered protein function. Increasing the nominal level reduces the number of calls, allowing a more selective subset to be retained when experimental capacity is limited. At the same time, an interval containing zero does not imply that the mutation has a negligible effect, and exclusion of zero does not by itself establish biological relevance or a functional consequence. When a minimum change of *δ >* 0 is required, the corresponding criteria are *L*_*i*_(*c*) *> δ* for candidate stabilization and *U*_*i*_(*c*) *<* −*δ* for candidate destabilization. These effect-size criteria were not evaluated here.

The conformal component is not intrinsically tied to ProbJanusDDG. Applying the same calibration procedure to a deliberately simple one-hot sequence predictor produced similar mutation-level calibration despite substantial differences in representation, architecture, and point-prediction performance. This illustrates the practical model-agnostic nature of the calibration strategy and suggests that the same approach could be used to equip other protein-stability predictors with calibrated uncertainty estimates. Overall, this work presents a protein-aware conformal strategy for complementing ΔΔ*G* point predictions with interpretable prediction intervals. ProbJanusDDG provides one competitive predictive instantiation of this strategy, including mutation-specific adaptive uncertainty, while the conformal calibration procedure remains transferable across contrasting predictors. Most importantly, the resulting intervals provide a practical selective criterion for prioritizing uncharacterized mutations with predicted stabilizing or destabilizing effects while explicitly identifying predictions for which the direction of the effect remains unresolved.

## 4 Materials and methods

### 4.2 Task and sign convention

For a wild-type protein sequence *x*_*w*_ and a mutant sequence *x*_*m*_, the model predicts ΔΔ*G*(*x*_*w*_, *x*_*m*_). We use the convention that positive ΔΔ*G* indicates stabilization and negative ΔΔ*G* indicates destabilization[17]. Reversing a mutation changes the sign of the physical target, since Gibbs free energy is a state function and the corresponding free-energy difference depends only on the initial and final states:

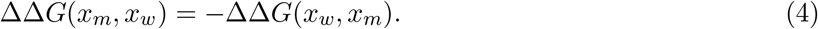

### 4.2 Datasets and splitting

The training set used in this work is S2450, a refined version of the widely used S2648 collection[8]. S2648 contains 2,648 experimentally characterized single-amino-acid substitutions across 131 proteins, with ΔΔ*G* values originally obtained from the ProTherm database[18]. In the construction of S2450, sequence similarity with the S669 benchmark was re-evaluated using full-length UniProt sequences rather than PDB sequences. Proteins from S2648 sharing more than 25% sequence identity with any protein in S669 were removed, resulting in the exclusion of 18 proteins and 198 mutations. The final S2450 dataset therefore contains 2,450 single-amino-acid substitutions from 115 protein sequences and provides a training set with reduced sequence overlap with the external benchmark.

For model development, we retained the five protein-disjoint benchmark folds associated with S2450. These folds were constructed using a 25% sequence-identity threshold to minimize sequence redundancy between training and validation subsets and contain 417, 772, 573, 511, and 177 mutations, respectively. Such protein-level separation is particularly important for ΔΔ*G* prediction, since multiple mutations are often available for the same protein and random mutation-level splitting could therefore lead to overly optimistic performance estimates[7].

External evaluation was performed on S669L and S461L, the full-length-sequence versions of two established blind benchmarks. S669[13] was specifically designed to assess generalization to proteins with low similarity to commonly used training collections, with sequence identity below 25% relative to datasets such as S2648 and VariBench[19]. The original benchmark contains both direct and reverse variants across 95 protein chains, with experimentally determined ΔΔ*G* values collected from ThermoMutDB and manually verified. In the present work, we use S669L, containing 669 mutations from 87 proteins after mapping to the corresponding full-length sequences. S461 is a curated benchmark derived from S669 and was introduced to remove possible problematic proteins [20]. It contains 461 single-site mutations with one experimental ΔΔ*G* measurement per variant. We use its full-length-sequence version, S461L, comprising 461 mutations from 46 proteins. Together, S669L and S461L provide complementary external tests of model generalization under low sequence similarity to the training data.

### 4.3 Model Architecture

In this section, we describe the architecture of the main model used in this study, illustrated in Figure 4a.

**Figure 4:**
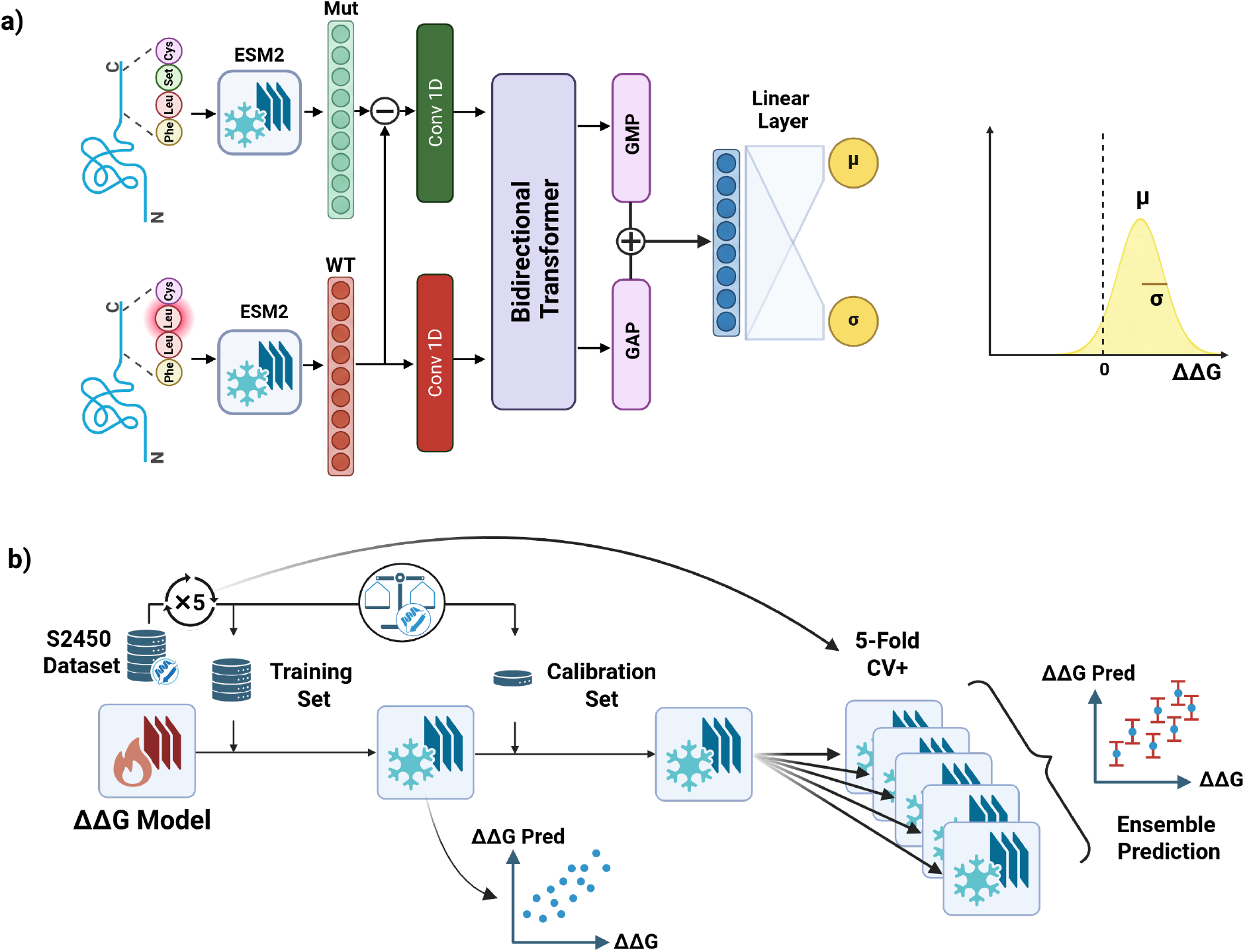
Schematic overview of ProbJanusDDG and the conformal calibration procedure. (a) Architecture of ProbJanusDDG, which jointly produces a central ΔΔ*G* prediction and a mutation-specific uncertainty estimate. (b) CV+ calibration pipeline used to combine fold-specific predictions and calibration residuals into standard and adaptive prediction intervals with coverage assessed empirically.

**Figure 5:**
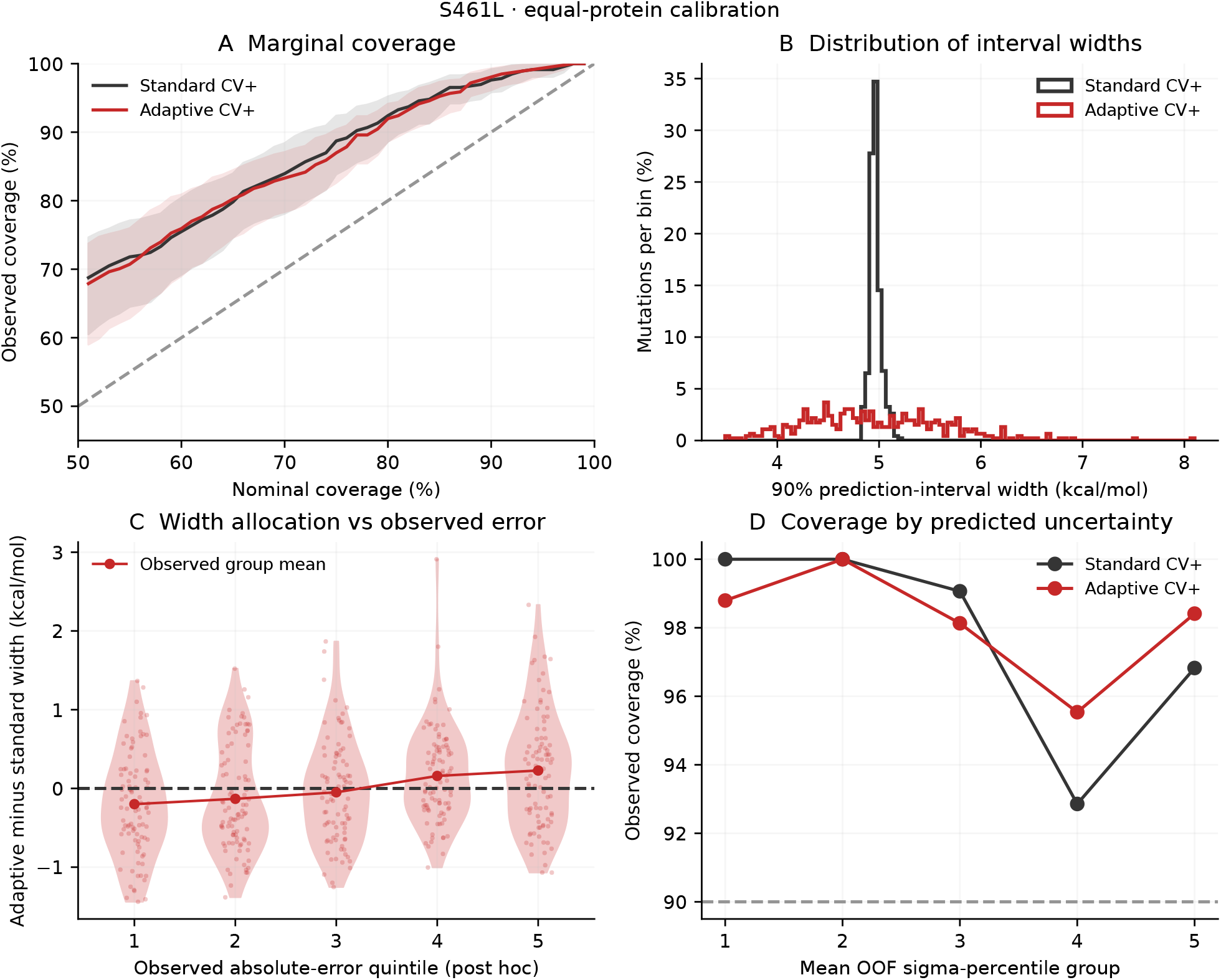
S461L counterpart of Fig. 1, using median predictions and protein-weighted calibration. Panels and bootstrap procedure are identical. S461L overlaps with S669L and is not an independent replication.

**Figure 6:**
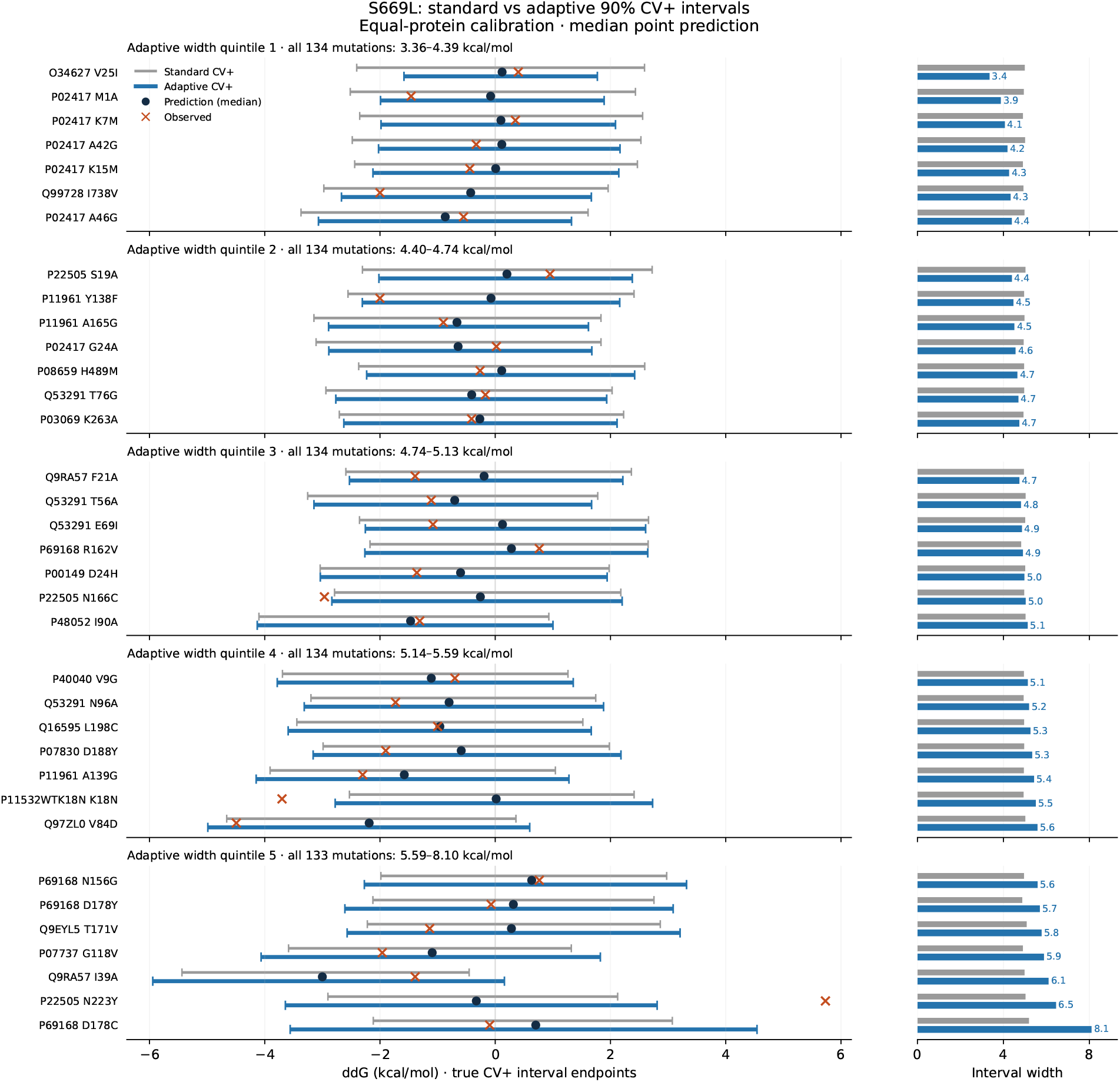
Width-stratified S669L examples with standard and adaptive protein-weighted 90% intervals and shared median predictions. Seven mutations are selected at evenly spaced width ranks within each of five adaptive-width quintiles, including each group’s endpoints. Selection uses interval width only, without target error or coverage. Gray and blue lines show actual standard and adaptive endpoints, slightly offset vertically for visibility; side bars compare their widths. The 35 examples are illustrative, not a separate test set.

**Figure 7:**
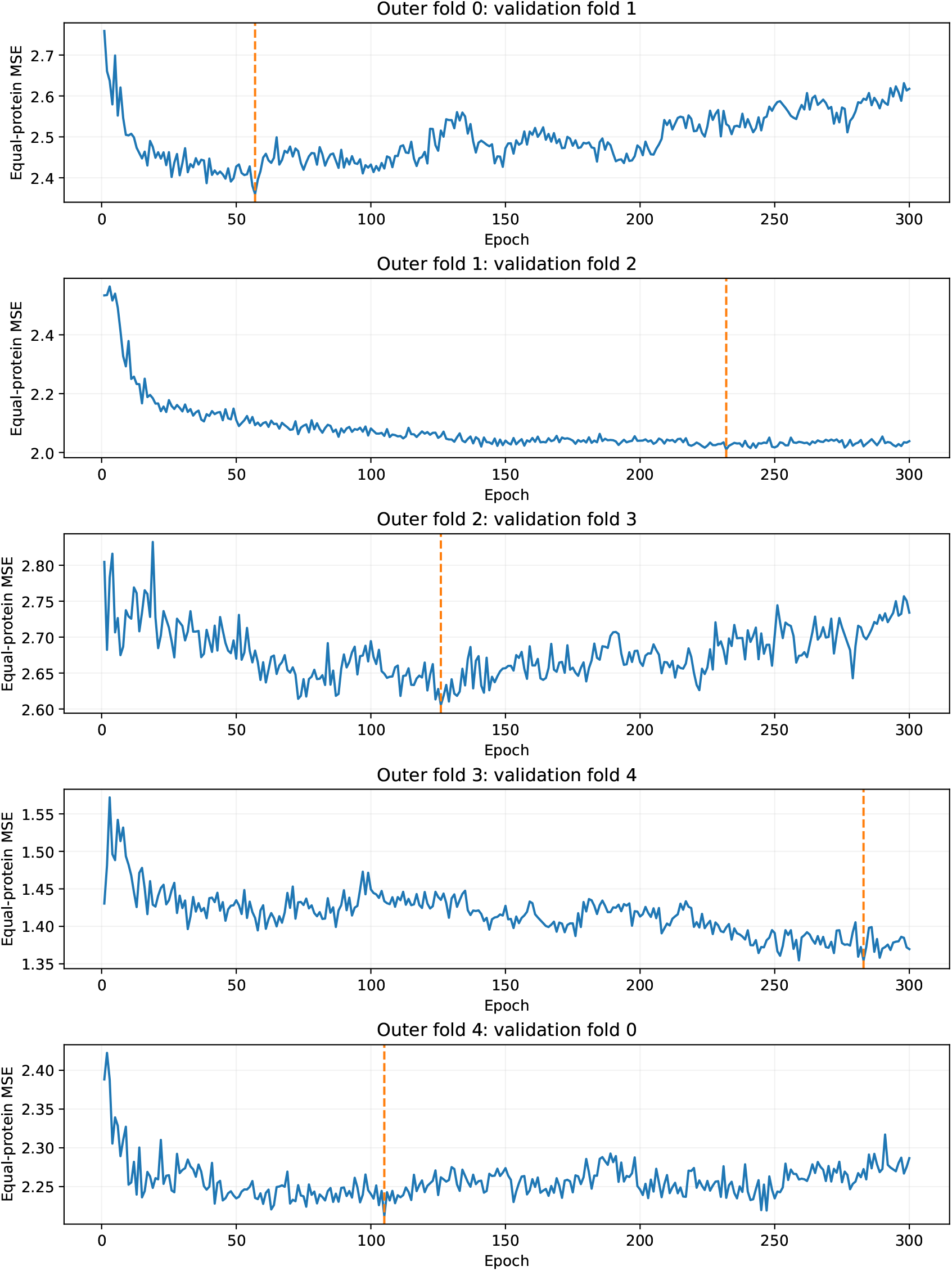
Protein-averaged MSE on the five internal validation folds over 300 epochs. Dashed lines mark the selected epochs 57, 232, 126, 283 and 105. Each curve comes from an inner model trained on three folds and excludes the associated outer calibration fold.

#### 4.3.1 Frozen protein language-model representations

Wild-type and mutant sequences were embedded independently using the 650-million-parameter ESM-2 model [21]. For each residue, ESM-2 provides a 1,280-dimensional contextual embedding. The language-model parameters were kept frozen throughout training, so ESM-2 was used only as a fixed feature extractor for the downstream ΔΔ*G* predictor.

#### 4.3.2 Bidirectional probabilistic architecture

The trainable network is based on the directional design of JanusDDG [7], with modifications to the output layer to support joint prediction of ΔΔ*G* and mutation-specific uncertainty. We deliberately retain this established backbone rather than introducing an additional sequence-processing architecture, allowing the present work to focus on the incorporation of mutation-specific uncertainty and conformal calibration. The original architecture is therefore adapted to produce both an antisymmetric ΔΔ*G* estimate and a symmetric uncertainty estimate, which is subsequently used to construct calibrated prediction intervals. A detailed description of the underlying sequence-processing architecture is provided in the original JanusDDG study.

In contrast to JanusDDG, which directly predicts a single ΔΔ*G* value, our model is designed to predict a probability distribution over ΔΔ*G*. Specifically, it employs two separate linear output heads to jointly estimate the parameters of a Gaussian distribution: the mean (*m*) and an unconstrained scale parameter (*s*). The predicted mean *m* is the directional central estimate, subsequently antisymmetrized within each fold model and aggregated across models. The scale parameter *s* is transformed through a softplus function to ensure positivity, yielding the standard deviation of the predicted Gaussian distribution:

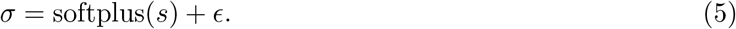

Thus, the model outputs both a point prediction, through *m*, and an associated estimate of predictive uncertainty, through *σ*.

At inference, the same trunk is evaluated in both the direct and inverse directions. If (*m*_*w→m*_, *s*_*w→m*_) and (*m*_*m→w*_, *s*_*m→w*_) denote the corresponding outputs, the delivered prediction is:

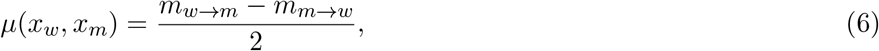

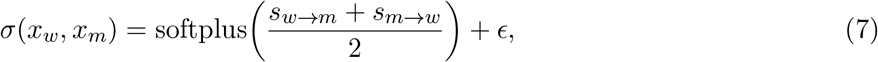

with *ϵ* = 10^−3^. Consequently, *µ*(*x*_*m*_, *x*_*w*_) = *−µ*(*x*_*w*_, *x*_*m*_) and *σ*(*x*_*m*_, *x*_*w*_) = *σ*(*x*_*w*_, *x*_*m*_) exactly.

### 4.4 Directional training with reverse augmentation

Within each fit, the network is optimized on the selected training mutations and their reverses, obtained by augmenting each observed mutation (*x*_*w*_, *x*_*m*_, *y*) with its reverse (*x*_*m*_, *x*_*w*_, −*y*). During training, the loss is applied directly to the single directional prediction produced for each sample; antisymmetrization is applied only at inference time.

Reverse augmentation encourages the directional predictor to learn the expected antisymmetric behavior of ΔΔ*G*. To illustrate this, any directional prediction *f* can be decomposed into an antisymmetric component *f*_*A*_ and a symmetric component *f*_*S*_, such that *f* = *f*_*A*_ + *f*_*S*_, where:

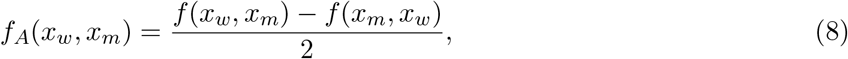

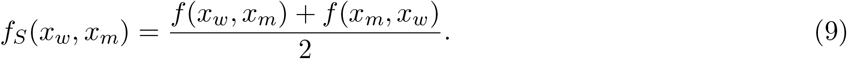

Considering the contributions of a mutation and its reverse to the overall squared-error objective gives:

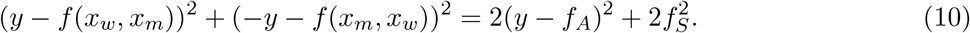

This identity illustrates the effect for an unweighted squared-error objective. The same mechanism also applies to the mean prediction in our Gaussian training objective, while the learned directional scales introduce additional terms in the loss.

At inference time, antisymmetry is enforced explicitly by evaluating the network in both directions and combining the two predictions as:

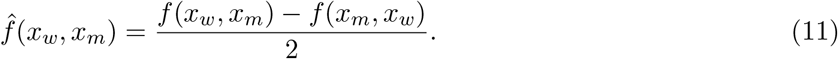

This operation extracts the antisymmetric component of the directional predictor and guarantees that reversing a mutation changes the sign of the predicted ΔΔ*G*. In contrast, if the loss is computed on an explicitly antisymmetrized prediction,

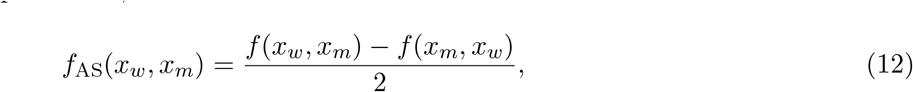

the symmetric component is removed by construction before the loss is evaluated. Therefore, no separate 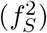 penalty arises: the optimization acts only on the antisymmetric prediction (*f*_AS_), which satisfies (*f*_AS_(*x*_*w*_, *x*_*m*_) = −*f*_AS_(*x*_*m*_, *x*_*w*_)) exactly.

The predictive mean and uncertainty are trained using the heteroscedastic Gaussian negative log-likelihood:

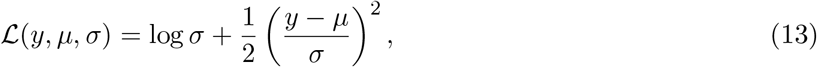

where the training outputs are directional and *σ* = softplus(*s*) + *ϵ*. This is the *β* = 0 member of the *β*-NLL family [22]. Optimization used Adam with a learning rate of 10^−4^, a batch size of 6 and a maximum of 300 epochs during cross-validation.

### 4.5 Nested epoch selection and four-fold refitting

Let *V*_0_, …, *V*_4_ denote the five fixed protein-disjoint folds. For outer fold *k*, the internal validation fold is *v*(*k*) = (*k* + 1) mod 5. A newly initialized model is trained for 300 epochs on the other three folds, excluding both *V*_*k*_ and *V*_*v*(*k*)_. The complete internal validation fold is used for epoch selection. If *P*_*v*_ is its set of WT proteins and *n*_*p,v*_ its number of mutations from protein *p*, the validation metric used for epoch selection is:

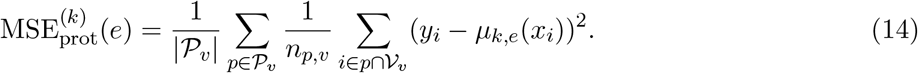

The prediction *µ*_*k,e*_ uses the antisymmetric inference rule. Each protein contributes equally to epoch selection, whereas the Gaussian training loss assigns equal weight to individual directional examples. This strategy keeps model optimization likelihood-based, while using a protein-balanced validation metric for epoch selection. By giving each protein equal weight, it reduces the influence of proteins represented by many mutations, although estimates for proteins with only a few variants may be less stable.

The selected epoch is the raw minimum over 1–300, with the earliest epoch resolving ties; no smoothing or patience is used. A new model is then initialized and fitted on all four folds other than *V*_*k*_ for that number of epochs, without validation. The number of epochs, rather than the number of optimizer steps, is transferred to the refit. Only this refitted model supplies the residuals on *V*_*k*_ and its external predictions. This construction ensures that, for each outer fold, calibration residuals are obtained from a model that has never used that fold during either training or model selection, thereby preserving a strict separation between model fitting and calibration. The five fitted models are retained; no full-S2450 final model is trained.

### 4.6 Protein-weighted empirical CV+ intervals

In this section, we describe how prediction intervals are constructed from the cross-validated model predictions and calibration residuals using a protein-weighted empirical CV+ procedure.

For mutation *i* in outer fold *k*(*i*), define:

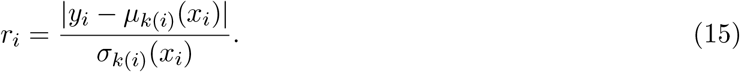

Each original mutation contributes once; reverse-augmentation duplicates are not calibration observations. For a new input *x*, the adaptive candidates are:

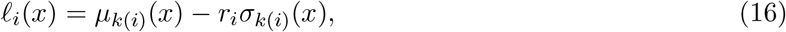

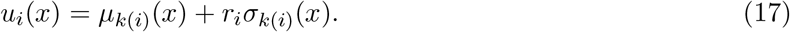

The standard candidates replace *r*_*i*_*σ*_*k*(*i*)_(*x*) with |*y*_*i*_ *− µ*_*k*(*i*)_(*x*_*i*_)|. The residual and the external prediction always use the same checkpoint. Multiplying every scale of a given model by the same positive constant cancels in its ratio of external to OOF scale, so no training-median scale normalization is needed.

If the calibration data contain *P* WT proteins and protein *p* has *n*_*p*_ mutations, observation *i* receives weight:

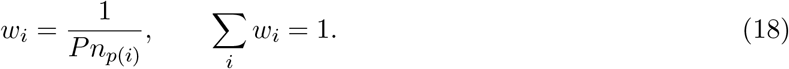

For candidate values *z*_*i*_, let

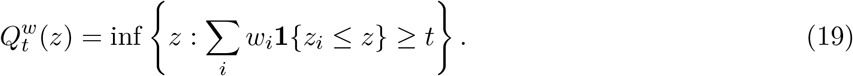

Our reported interval is:

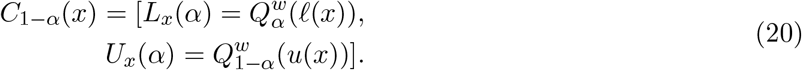

We refer to 1 − *α* as the nominal level of this empirical construction, not as a proven lower bound on its coverage.

Supplementary section A.3 provides finite-sample coverage bounds for this protein-balanced construction under aligned hierarchical sampling and data-independent protein-level fold allocation. For the empirical quantile rule used here, marginal coverage is bounded below by 1 − 2*α* − *ε*_*P,K*_(*α*), where *P* is the number of calibration proteins and the finite-sample correction is specified in the appendix. Under the additional assumptions of balanced folds and a symmetric fitting procedure, this correction is at most 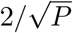, yielding an asymptotic lower bound of 1 − 2*α*. These assumptions are not established by the present sequence-based partitioning and external benchmark curation; coverage under the experimental protocol is therefore evaluated empirically.

The reported point prediction is

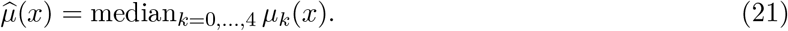

It is exactly antisymmetric because it is the median of five antisymmetric predictions. The interval endpoints are computed from candidates, not by centering a pooled residual quantile on this median. Consequently, standard CV+ widths are not exactly constant and neither interval rule is required to be symmetric around 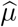.

### 4.7 Assumptions and scope of validity

The proposed calibration framework reflects the hierarchical structure of the data, in which multiple mutations are observed for each protein and proteins contribute unequal numbers of measurements. Its interpretation therefore depends on the target sampling population, the separation between model fitting and calibration, and the transferability of the learned predictive scale to unseen proteins.

#### Protein-balanced target

Calibration assigns the same total weight to each protein, irrespective of the number of observed mutations. This corresponds to a two-stage target population in which a protein is sampled first and a mutation is then sampled from that protein. Weighting mutations from protein *g* proportionally to 1*/n*_*g*_ prevents densely sampled proteins from dominating the calibration distribution and shifts the target from mutation-level to protein-balanced performance.

We treat the full wild-type sequence as the operational protein-level cluster. The intended interpretation is that calibration proteins are representative of the population of proteins encountered at deployment, with mutations sampled according to comparable within-protein mechanisms. The weighting corrects unequal representation across observed proteins, but is not intended to correct arbitrary selection bias or distribution shift.

#### Out-of-fold separation

All calibration scores are obtained from models that exclude the corresponding outer-fold protein. The same fold-specific model is then used to generate the prediction-time candidate associated with that score, preserving the CV+ score–prediction pairing. Model selection is performed within the training portion of each outer split, and the held-out protein is therefore excluded from both parameter estimation and epoch selection.

Outer-fold models are trained on overlapping subsets of the data, so the resulting scores are not mutually independent. This dependence is intrinsic to cross-validation and does not alter the fact that each score is computed out of fold.

#### Learned predictive scale

Calibration uses the normalized score

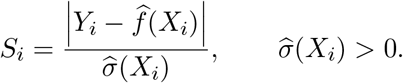

The same prediction and scale functions are used for out-of-fold observations and new inputs. No Gaussian residual assumption is required: 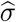 acts as a learned local scaling factor that allows prediction intervals to adapt to heterogeneous prediction difficulty.

Useful adaptivity requires the relationship between predicted scale and realized error to transfer to unseen proteins. The resulting intervals should therefore be interpreted as prediction intervals for future measured ΔΔ*G* values, including measurement variability represented in the training data.

#### Coverage interpretation

The method retains the model-specific construction of CV+ while replacing mutation-uniform aggregation with protein-balanced weighted quantiles. Weighted conformal procedures can retain finite-sample guarantees when the weighting scheme is matched to the target sampling distribution; our weights,

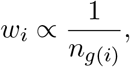

define a protein-balanced empirical target. We therefore evaluate calibration primarily through empirical coverage on unseen proteins, together with interval width and scale-calibration diagnostics. The method is designed to provide calibrated uncertainty at the protein-population level; it is not intended to guarantee identical coverage for every individual protein, mutation, predicted scale, or ΔΔ*G* subgroup.

### 4.8 Evaluation

Evaluation was performed separately on S669L and S461L using the five fitted models and the calibration residuals obtained from S2450. External observations were used exclusively for evaluation. Because S461L is a curated subset overlapping S669L, its results were interpreted as an assessment under a different benchmark composition rather than an independent replication. Point-prediction performance was assessed using Pearson and Spearman correlations, mean absolute error (MAE), and mean squared error (MSE), calculated for the median ensemble prediction with equal weight assigned to each test mutation.

Prediction intervals were evaluated through empirical coverage, width, and interval score. Standard and adaptive CV+ used the same fitted models, out-of-fold observations, and equal-protein calibration weights, allowing their comparison to isolate the effect of residual normalization by the learned scale. For a test set containing *N* mutations from *P*_test_ distinct full-length WT sequences, we calculated both mutation-level and protein-averaged coverage:

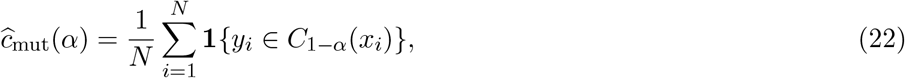

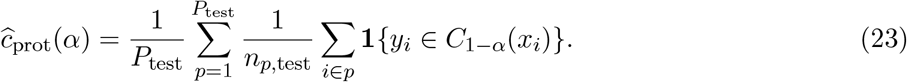

The latter gives each protein equal influence and is aligned with the protein-balanced calibration target. Coverage curves were evaluated at integer nominal levels from 51% to 99%, and calibration discrepancy was summarized by the mean absolute difference between mutation-level coverage and the nominal level across this grid.

Detailed interval comparisons were performed at the 90% nominal level. We reported the mean full interval width, *u*_*i*_ *− l*_*i*_, and the mean interval score,

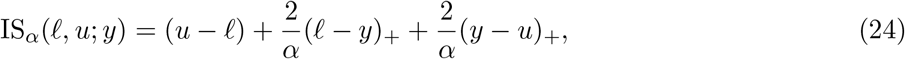

where (*a*)_+_ = max(*a*, 0). Lower scores indicate a better balance between interval width and penalties for uncovered observations. Width and interval score were averaged with equal weight per mutation and were used only for evaluation, not model selection.

To assess the information carried by the learned scale, each model’s scale for a test mutation was converted to its percentile within that model’s held-out calibration-scale distribution, using midranks for ties. The mean of the five percentiles defined an uncertainty rank, whose association with absolute median-prediction error was measured using Spearman correlation. Fixed percentile boundaries at 0.2, 0.4, 0.6, and 0.8 defined five uncertainty groups, which need not contain equal numbers of test mutations. Coverage heterogeneity was summarized by the population standard deviation of the five mutation-level group coverages, expressed in percentage points.

The redistribution of interval width was examined through paired adaptive-minus-standard width differences, including summaries within quintiles of observed absolute prediction error. These error-based groups were used exclusively for post-hoc interpretation. All mutations whose coverage status changed between the two interval definitions were displayed as illustrative cases.

The practical utility of adaptive intervals was assessed through directional calls at integer nominal levels from 50% to 95%. Under our sign convention, an interval lying strictly above zero indicated a stabilizing effect, whereas an interval lying strictly below zero indicated a destabilizing effect; intervals containing or touching zero yielded no directional call. At each level, we reported the proportion of all test mutations receiving a call, the observed sign accuracy among called mutations, and the proportion of distinct WT proteins with at least one call. Sign accuracy assigned equal weight to each called mutation; observed targets equal to zero counted as incorrect calls, and accuracy was undefined when no calls were made. These empirical selective accuracies were distinguished from nominal interval coverage.

Where reported, 95% uncertainty intervals were obtained from 4,000 paired protein-cluster bootstrap resamples, retaining all mutations of each sampled protein and using the same resamples for standard and adaptive intervals. Fitted models, calibration residuals, and diagnostic group boundaries were held fixed. Coverage bands were pointwise; bootstrap intervals therefore describe variability across sampled test proteins without incorporating model-fitting variability.

### 4.9 Naive sequence-based baseline model

To assess whether the proposed uncertainty-quantification procedure is independent of the underlying predictive architecture, we implemented a deliberately simple sequence-based baseline model. The model operates directly on the full-length wild-type and mutant protein sequences and does not use pretrained protein language models, attention mechanisms, convolutional layers, or any form of external pretraining.

Wild-type and mutant residues are represented using a 20-dimensional one-hot encoding. For each residue position *i*, the model receives a 42-dimensional feature vector

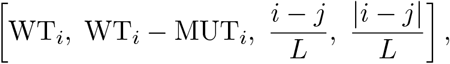

where WT_*i*_ and MUT_*i*_ denote the one-hot representations of the wild-type and mutant residues, *j* is the position of the single amino-acid substitution, and *L* is the sequence length. The mutation site *j* is identified directly by comparing the wild-type and mutant sequences. The two positional terms encode the signed and absolute distance from the mutated residue, normalized by sequence length.

Each residue is processed independently by the same shared feed-forward neural network,

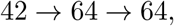

with ReLU activations after each linear layer. The resulting per-residue representations are then aggregated using three complementary summaries: the mean across valid residues, the element-wise maximum across valid residues, and the 64-dimensional representation corresponding to the mutated site. Their concatenation yields a 192-dimensional protein-level representation. Padding positions are always excluded from the aggregation, allowing sequences of arbitrary length to be processed without imposing a fixed maximum sequence length.

The aggregated representation is passed to a final readout network,

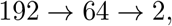

again using a ReLU activation after the first linear layer. The two outputs correspond to a directional point estimate *m*_*AB*_ and an unconstrained scale parameter *s*_*AB*_. The complete model contains 19,394 trainable parameters.

Training is performed directionally. For a mutation from sequence *A* to sequence *B*, the predicted mean and scale are defined as

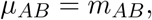

and

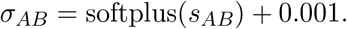

The training set is augmented with the corresponding reverse mutations, for which the sign of the experimental ΔΔ*G* label is inverted. This exposes the model explicitly to both mutation directions during optimization.

At inference time, predictions from the forward and reverse directions are combined to enforce the desired symmetry properties. The final point estimate is obtained as

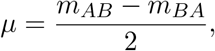

which is exactly antisymmetric under reversal of the mutation. The uncertainty scale is instead computed as

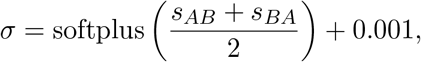

yielding an exactly symmetric uncertainty estimate for the direct and reverse mutation pair.

Because residues are processed independently before global aggregation, the model has no explicit mechanism for learning long-range residue interactions. Its representation and architecture therefore differ substantially from those of the main predictor. This intentionally low-capacity baseline provides a stringent test of model agnosticism: if conformal calibration remains effective despite these architectural and representational differences, its validity is less likely to depend on the specific structure of the underlying ΔΔ*G* predictor.

## 4.10 Code availability

The implementation and scripts used for training, calibration, evaluation, and figure generation are available at https://github.com/bardu98/ProbJanusDDG.

## 5 AI Disclosure Statement

Generative artificial intelligence tools were used during manuscript preparation to improve the clarity, grammar, and overall presentation of the text, and to assist with code cleaning and refactoring. All scientific content, analyses, results, interpretations, and final code were reviewed and validated by the authors, who takes full responsibility for the work.

## A Supplementary

### A.1 Method performance on external datasets

To place the predictive performance of ProbJanusDDG in the context of existing ΔΔ*G* predictors, we compared its point-prediction accuracy with that of recent state-of-the-art methods on the external S669L and S461L benchmarks. The comparison includes both sequence-based and structure-based approaches, allowing ProbJanusDDG to be evaluated against methods that differ substantially in input representation, architecture, and use of structural information.

Performance was assessed using the same external test sets adopted in the main analysis. For ProbJanusDDG, predictions correspond to the median ensemble output across the five refitted models. The benchmark metrics considered were Pearson correlation and mean absolute error (MAE), whenever the corresponding values were available for the reference methods. The complete comparison is reported in Supplementary Table 4.

On S669L, ProbJanusDDG achieved predictive performance comparable to the strongest methods reported for this low-homology benchmark, indicating that the introduction of uncertainty estimation and conformal calibration does not come at the expense of competitive point-prediction accuracy. On S461L, performance was higher across the main regression metrics, further supporting the ability of the model to generalize to an independently curated external set.

Overall, these results show that ProbJanusDDG remains competitive with current state-of-the-art protein-stability predictors while additionally providing mutation-specific uncertainty estimates and calibrated prediction intervals, which are not available from most conventional ΔΔ*G* prediction methods.

### A.2 Direction calls at all nominal levels

Table 5 summarizes the results at selected nominal coverage levels between 50% and 90%, providing the numerical values corresponding to Fig. 3.

### A.3 Protein-balanced CV+ prediction intervals and finite-sample coverage

Multiple mutations measured on the same protein form a sampling cluster. Consequently, equal weighting of mutations gives greater calibration influence to proteins represented by more measurements, and ordinary observation-level exchangeability need not hold. We therefore formulate calibration for a target population in which a protein is sampled first and a mutation is then sampled within that protein. The resulting prediction intervals preserve the association between each out-of-fold score and its fold-specific predictor, while assigning equal total calibration mass to each protein. This construction connects CV+ [16, 23] with hierarchical conformal inference [24]. We distinguish the empirical quantile rule used in our experiments from a finite-sample-corrected rule and establish marginal coverage bounds under explicit sampling and fitting assumptions. All results below are lower bounds on marginal coverage, rather than assertions of exact nominal coverage.

#### A.3.1 Hierarchical sampling and the prediction target

Let *P* denote the number of observed proteins, and write

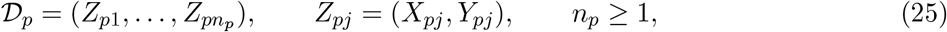

where *X*_*pj*_ denotes the mutation input and *Y*_*pj*_ its measured ΔΔ*G* value. The following sampling model provides sufficient conditions for the guarantees.

##### Assumption 1

(Hierarchical sampling and target alignment). *Protein-level latent variables* Θ_1_, …, Θ_*P*_ *are independent draws from a common distribution Q. Conditional on* Θ_*p*_, *the observed cluster size is drawn from a distribution on the positive integers, which may depend on* Θ_*p*_. *Conditional on* (Θ_*p*_, *n*_*p*_),

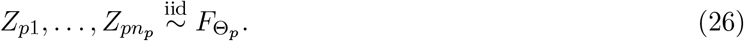

*Clusters are independent across proteins. The test observation is generated independently by*

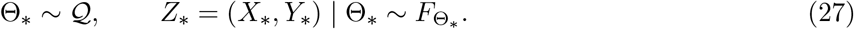

*In particular, the within-protein observation law in* (26) *does not change with n*_*p*_ *after conditioning on* Θ_*p*_.

The protein-balanced target law is thus

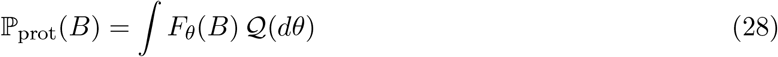

for measurable *B*. Equal protein weighting does not require *Q* to be uniform over all possible protein sequences. Rather, it prevents the number of measured mutations from introducing an additional size bias into the target mixture. Its empirical counterpart is

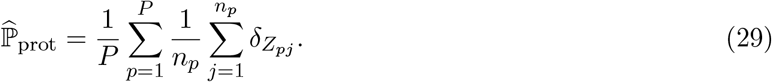

For any fixed bounded measurable function *h*, this empirical measure satisfies 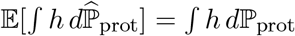. This identity identifies the sampling target; it does not, by itself, establish validity for data-dependent prediction intervals.

Assumption 1 allows arbitrary between-protein heterogeneity and unequal cluster sizes. Within-protein independence is a sufficient condition, rather than a necessary feature of the argument. The proofs also apply under hierarchical exchangeability of the observed and augmented test clusters, provided a uniformly selected observation from the test cluster has the target law. For the augmented-fold argument below, exchangeability must extend to all *P* + *P/K* clusters involved. The hierarchical independent sampling model supplies this extension automatically.

#### A.3.2 Fold-specific scores and weighted endpoint aggregation

Partition the protein indices into *K* nonempty folds *I*_1_, …, *I*_*K*_, with *P*_*k*_ = |*I*_*k*_| and ∑_*k*_ *P*_*k*_ = *P*. Let *k*(*p*) denote the fold containing protein *p*.

##### Assumption 2

(Fold allocation and out-of-fold fitting). *Fold allocation is independent of the protein data, with prescribed fold sizes. For each k, the fitted functions* (*µ*_*k*_, *σ*_*k*_), *including any preprocessing, tuning, or model selection, depend only on clusters outside I*_*k*_ *and on auxiliary randomness independent of the calibration and test data. The scale satisfies* 0 *< σ*_*k*_(*x*) *< ∞, and predictions are finite at the inputs considered*.

Define the normalized score and out-of-fold residual by

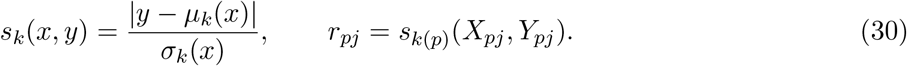

For a new input *x*, the candidate endpoints are

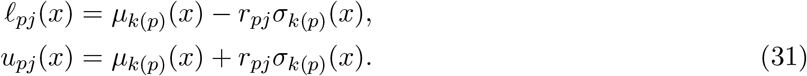

The same fitted model generates both *r*_*pj*_ and its test-time endpoints. Only original measured mutations enter calibration; reverse augmentation does not create additional calibration observations. The unnormalized construction is recovered by setting *σ*_*k*_ ≡ 1. Neither construction requires Gaussian residuals or a correctly specified predictive scale.

Each mutation receives weight

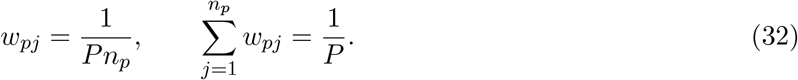

For a collection *v* = (*v*_*pj*_), define

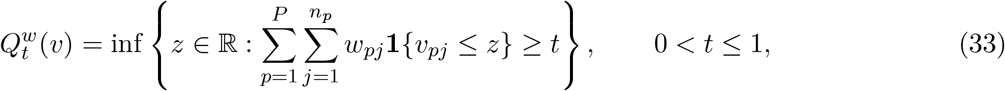

with *Q*^*w*^(*v*) = +*∞* for *t >* 1. For 0 *< α <* 1*/*2, the empirical interval is

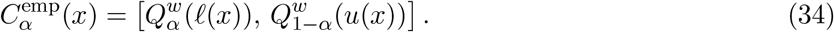

We also define a corrected interval using

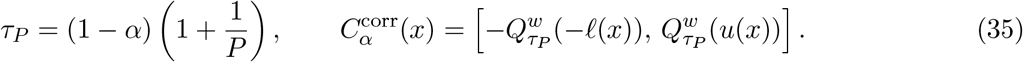

The reflected lower quantile specifies the endpoint convention in the presence of atoms or ties. The correction is equivalent to assigning mass 1*/*((*P* + 1)*n*_*p*_) to each observed candidate and mass 1*/*(*P* + 1) to the unobserved test protein, represented conservatively at for the lower endpoint −∞ and +∞ for the upper endpoint. If *τ*_*P*_ *>* 1, the corrected interval is ℝ. This is a protein-level correction: the relevant count is *P*, rather than ∑_*p*_ *n*_*p*_.

#### A.3.3 Coverage bounds from hierarchical foldwise ranks

##### Proposition 1

(Coverage with possibly unequal fold sizes). *Under Assumptions 1 and 2*,

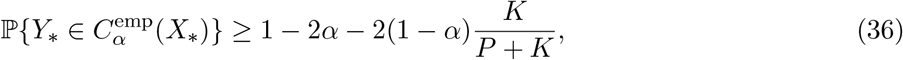

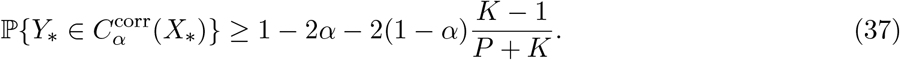

*The bounds are marginal over the development sample, independent fold allocation, fitting randomness, and test observation. Fold sizes need not be equal, and the fitted models need not be independent*.

*Proof*. For each fold, define

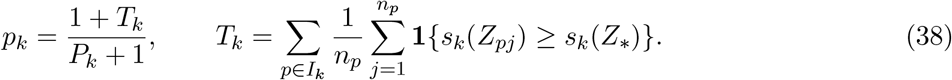

We first establish ℙ(*p*_*k*_ ≤ *t*) ≤ *t* for 0 ≤ *t* ≤ 1. Condition on the training clusters outside *I*_*k*_ and the fitting randomness, so that *s*_*k*_ is fixed. Augment the calibration data by an unobserved test cluster generated using the same sampling mechanism, and select the test observation uniformly within this cluster. This produces the target distribution in (27).

For a realized collection of *P*_*k*_ + 1 clusters, give each observation *a* = (*p, j*) weight *b*_*a*_ = 1*/*((*P*_*k*_ + 1)*n*_*p*_) and define its full upper-tail rank

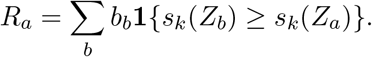

The deterministic weighted-rank inequality ∑_*a*_ *b*_*a*_**1 {***R*_*a*_ ≤ *t*} ≤ *t* holds, including with tied scores. Replacing the contribution from the observation’s own cluster by its maximum possible mass 1*/*(*P*_*k*_ + 1) yields a value at least *R*_*a*_. For an observation in the test cluster, this conservative value is precisely *p*_*k*_. Exchangeability of the *P*_*k*_ + 1 clusters and uniform selection within the test cluster therefore give ℙ(*p*_k_ ≤ *t*) ≤ *t*.

Set *T* = ∑_*k*_ *T*_*k*_. If *y* is above the empirical upper endpoint, the weight of candidates satisfying *u*_*pj*_(*x*) *< y* is at least 1 *− α*. If *y* is below the empirical lower endpoint, the weight satisfying *l*_*pj*_(*x*) *> y* is greater than 1 *− α*. In either case, each such candidate satisfies *s*_*k*(*p*)_(*x, y*) *> r*_*pj*_. Consequently,

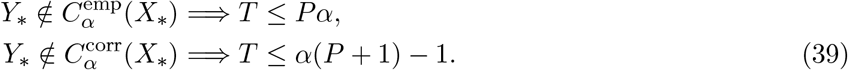

For the second implication, the mass of excluding candidates is at least *τ*_*P*_; if *τ*_*P*_ *>* 1, noncoverage is impossible.

The convex combination

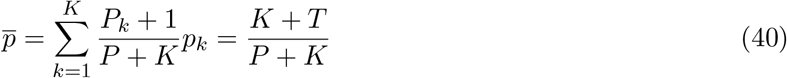

satisfies 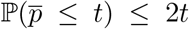, without independence assumptions. To verify this, for 0 *< t* ≤ 1*/*2 let *a*_*k*_ = (*P*_*k*_ + 1)*/*(*P* + *K*) and note that

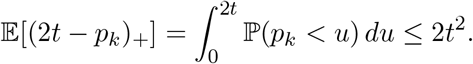

On 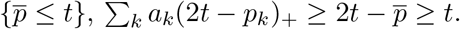. Markov’s inequality proves the claim; for *t >* 1*/*2 it follows from the trivial probability bound. Substituting the two thresholds in (39) into (40) gives

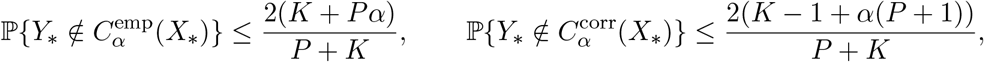

which are the stated bounds.

#### A.3.4 Sharper bounds under symmetric fitting and balanced folds

The preceding proposition requires only conditional alignment of each held-out fold with the test population. A second bound can be obtained by a weighted tournament argument, but it requires stronger symmetry of the complete fitting procedure.

##### Assumption 3

(Balanced folds and a symmetric fitting rule). *K divides P, and each fold contains m* = *P/K proteins. All fitted prediction and scale functions are outputs of the same measurable fitting rule A, which is invariant to permutations of training proteins and to permutations of observations within each protein. This invariance includes preprocessing and model selection. A sufficient randomized implementation uses a shared data-independent random seed for all evaluations of A, with the stated invariance holding conditional on that seed. More general randomized implementations require equivariance of the joint auxiliary fitting construction used in the proof*.

##### Lemma 1

(Weighted tournament inequality). *Let* {*b*}_*a*_ *be nonnegative weights summing to one, and let A*_*ab*_ ∈ {0, 1} *satisfy A*_*ab*_ + *A*_*ba*_ ≤ 1. *Set W*_*a*_ = ∑_*b*_ *b*_*b*_*A*_*ab*_. *Then, for* 0 ≤ *t* ≤ 1,

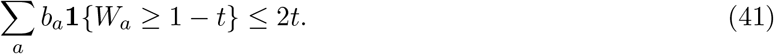

*Proof*. Let *S* = {*a* : *W*_*a*_ ≥ 1 − *t*} and *B* = ∑_*a*∈*S*_ *b*_*a*_. The antisymmetry condition bounds the contribution of comparisons within *S* by *B*^2^*/*2, while comparisons with its complement contribute at most *B*(1 *− B*). Thus

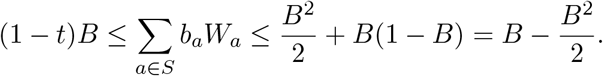

If *B >* 0, division by *B* gives *B* ≤ 2*t*; the result is immediate for *B* = 0.

##### Theorem 1

(Protein-balanced CV+ coverage). *Under Assumptions 1, 2, and 3*,

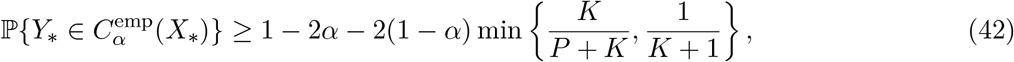

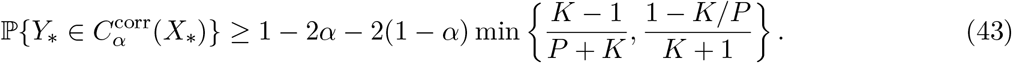

*Proof*. Augment the *P* development proteins with *m* independent test proteins, producing *M* = *P* + *m* proteins in *K* + 1 folds of size *m*. For two distinct folds *k* and *l*, fit *A* using all clusters outside *I*_*k*_ ∪ *I*_*l*_, and denote the resulting normalized score by *s*_−(*k,l*)_. For observation indices *a* = (*p, j*) and *b* = (*q, h*), define

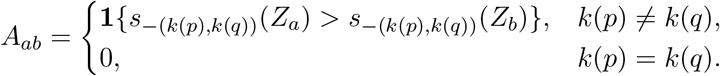

Both observations are evaluated using the same model, so *A*_*ab*_ + *A*_*ab*_ *≤* 1. Assign *b*_pj_ = 1*/*(*Mn*_p_), and put *W*_*a*_ = ∑_*b*_ *b*_*b*_*A*_*ab*_. The symmetric fitting rule makes the construction equivariant under permutations within folds and permutations of whole folds. Together with hierarchical sampling, this implies that the probability of *W*_*a*_ ≥ *c* for a uniformly selected observation from any specified protein equals *E*[∑_*a*_ *b*_*a*_ {*W*_*a*_ ≥ *c*}].

Take a protein in the added test fold. Removing its fold and a development fold leaves exactly the data used by the corresponding original out-of-fold model. The endpoint argument in (39) therefore yields

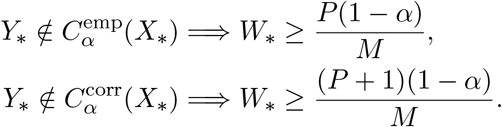

The corrected interval is ℝ when *τ*_*P*_ *>* 1; hence its bound is automatic in that case. Otherwise both thresholds lie in [0, 1]. Lemma 1 gives

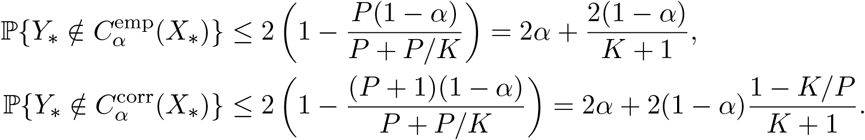

Combining these bounds with Proposition 1 proves the theorem.

##### Corollary 1

(A fold-independent lower bound). *Under the assumptions of Theorem 1, either interval construction satisfies*

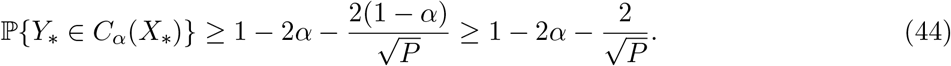

*Proof*. If 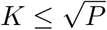, then 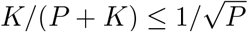 otherwise 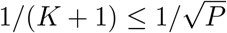. Thus the minimum in (42) is at most 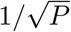. Each corresponding term in (43) is no larger, giving the same conclusion.

The corrected bound has the same structure as the classical CV+ bound, since

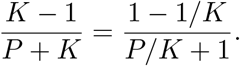

The substitution of *P* for the mutation count is justified by the hierarchical rank and tournament arguments, rather than by an assumed scalar effective sample size. In the leave-one-protein-out case *K* = *P*, the corrected guarantee reduces to 1 − 2*α*, whereas the stated empirical-rule bound retains a finite correction of 2(1 − *α*)*/*(*P* + 1). Increasing the number of mutations within an already observed protein can improve score estimation and model fitting, but does not increase the number of protein-level sampling units in these bounds.

#### A.3.5 Interpretation and applicability to the experimental protocol

These results concern a mutation sampled from a new protein drawn from the same protein population as the development clusters. Equivalently, they bound the expectation, over the development data and a new protein, of that protein’s mutation-level coverage under its within-protein sampling law. They do not ensure identical coverage for every protein, simultaneous coverage of all mutations in a protein, coverage conditional on a fixed development sample, or validity under an arbitrary change in the mutation-selection mechanism.

The parameter 1 − *α* is the nominal level of the quantile construction. The distribution-free lower bounds involve 1 − 2*α* and finite-sample corrections; in particular, *α* = 0.10 does not yield a theorem guaranteeing 90% coverage. As an illustration under the theorem’s assumptions, *P* = 115, *K* = 5, and *α* = 0.10 give lower bounds of 0.725 for the empirical rule and 0.740 for the corrected rule. These values describe hypothetical aligned sampling and eligible folds, not guaranteed coverage on the external benchmarks.

In the present experiments, the empirical rule (34) is used with fixed protein-disjoint folds constructed to reduce sequence similarity between training and validation subsets. Such separation prevents direct reuse of a held-out protein during fitting, but does not establish the data-independent fold allocation in Assumption 2. In addition, sequence-based filtering and curation of external benchmarks do not establish that their proteins and mutations follow the same hierarchical sampling law as the development data. The prescribed cyclic choice of an internal validation fold establishes outer-fold exclusion, but does not by itself verify the symmetry of the complete fitting rule required in Assumption 3.

Accordingly, the theorems provide a theoretical justification for protein-balanced CV+ under explicitly aligned hierarchical sampling and fitting conditions. They are not asserted as finite-sample guarantees for the current low-homology benchmark protocol. Coverage on those benchmarks is assessed empirically, with protein-averaged coverage reporting the metric aligned with the intended target and mutation-averaged coverage reporting a different weighting of the evaluated observations.

## B Supplementary figures

